# Green genetically encoded IP_3_ biosensor for hierarchical analysis of its signaling network

**DOI:** 10.64898/2026.05.12.724711

**Authors:** Li Tian, Keisuke Yamashita, Zixu Feng, Takashi Tsuboi, Takanobu Yasuda, Bo Zhu, Tetsuya Kitaguchi

## Abstract

Inositol 1,4,5-trisphosphate (IP_3_) is a key second messenger that regulates diverse physiological processes. Visualization of IP_3_ dynamics in living cells is therefore important for understanding its signaling processes. In this study, we developed genetically encoded green fluorescent IP_3_ biosensors named Green iPenguins with distinct half-maximal effective concentrations (EC_50_) for IP_3_, enabling detection of IP_3_ signals over a range of concentrations. The biosensors displayed more than a 4-fold increase in fluorescence intensity upon IP_3_ and showed high specificity for IP_3_ over structurally related molecules. When expressed in HEK293T cells, the biosensors enabled visualization of IP_3_ dynamics involved in different signaling pathways. They were also compatible with dual-color imaging, allowing simultaneous monitoring of IP_3_ together with cAMP or Ca^2+^ signals. In addition, the hierarchical relationship between IP_3_ and Ca^2+^ signaling was visualized, providing insight into the temporal relationship between these two second messengers. The biosensors are expected to facilitate future studies of physiological processes involving IP_3_ signaling networks.

## 1 Introduction

Inositol 1,4,5-trisphosphate (IP_3_) is a second messenger that is involved in a wide range of physiological processes, including apoptosis^1^, secretion^2^, gene expression^3^, fertilization^4^, and neuronal activity^5^. Upon stimulation of G protein-coupled receptors (GPCRs) that couple to Gq protein, phospholipase C (PLC) is activated, leading to IP_3_ production at the plasma membrane. The IP_3_ subsequently diffuses through the cytosol to activate IP_3_ receptors on the endoplasmic reticulum, triggering Ca^2+^ mobilization^6^. GPCR-mediated IP_3_ signaling serves as a key regulator of various cellular responses through Ca^2+^ signaling, while GPCRs also activate a variety of downstream signaling pathways^7^, including cAMP, small G proteins, cGMP, etc., highlighting the importance of analyzing diverse signaling molecules for a comprehensive understanding of physiological processes. To date, a variety of genetically encoded biosensors have been developed, providing a powerful approach for visualizing the dynamics of the intracellular molecules in living cells with high spatiotemporal resolution.

GFP fused to the PH domain of PLCδ, GFP-PHD^8^ was the first fluorescent probe used to monitor intracellular IP_3_, as IP_3_ displaces it from PIP_2_ at the plasma membrane. However, its specificity is limited due to PIP_2_ dependence, and monitoring probe release from the membrane provides only an indirect readout, motivating the exploration of alternative approaches. For direct visualization of IP_3_, the genetically encoded IP_3_ biosensor was developed based on Förster resonance energy transfer (FRET), and multiple FRET-based biosensors were subsequently engineered. The first-generation FRET-based IP_3_ biosensor LIBRA^9^ inserted the ligand-binding domain of IP_3_ receptor between fluorescent proteins, CFP and YFP, enabling ratiometric measurement of intracellular IP_3_ dynamics in living cells. Subsequently, several FRET biosensors, such as Fretino^10^ revealed synchronous IP_3_ dynamics in neuronal dendrites. The biosensors^11–13^, including the IRIS series, were used to investigate their relationship with Ca^2+^ signaling at a single cells, and additional biosensors^14–16^ were further improved for user friendliness.

As mentioned above, IP_3_ dynamics have been imaged with Ca^2+^ dynamics using fluorescent chemical dyes or genetically encoded biosensors at a single cells. Since this imaging requires detection of at least two emission channels for the FRET-based IP_3_ biosensor, along with one or sometimes two additional channels to monitor other signaling molecules, the microscope setup becomes more complex, which has limited its use in multi-color imaging. In contrast, single fluorescent protein (FP)-based biosensors are well suited for multi-color imaging because each biosensor requires only a single emission, making simultaneous monitoring of multiple signaling molecules simple and straightforward. However, the absence of single FP-based biosensors for IP_3_ has hindered simultaneous monitoring, thereby limiting analysis of signaling network dynamics. Consequently, there has been a strong need for IP_3_ biosensor compatible with multi-color imaging.

Here, we developed single FP-based IP_3_ biosensors, named as Green iPenguins (<u>Green</u> IP_3_ biosensor based o<u>n</u> sin<u>g</u>le fl<u>u</u>orescent prote<u>in</u>), and performed live-cell imaging. By simplifying multi-color imaging, we anticipate that Green iPenguins will facilitate studies of IP_3_ -dependent physiological processes that have been difficult to address with existing biosensors.

## 2 Materials and methods

### 2.1 Chemicals

Inositol 1,4,5-trisphosphate (IP_3_(, inositol 1,3,4-trisphosphate (1,3,4-IP_3_(, inositol 1,4-bisphosphate (1,4-IP_2_), inositol 1,3,4,5-tetrakisphosphate (1,3,4,5-IP_4_), phosphatidylinositol 4,5-bisphosphate (PIP_2_), diacylglycerol (DAG), and ionomycin were purchased from Cayman Chemical (Ann Arbor, MI, USA). Charbacol, histamine, and adrenalin were purchased from Merck KGaA (Darmstadt, Germany). Adenosine triphosphate (ATP) was purchased from Oriental Yeast Co., Ltd. (Tokyo, Japan). U73122 was purchased from FUJIFILM Wako Pure Chemical Corporation (Osaka, Japan)

### 2.2 Plasmid constructions

The cDNA encoding the IP_3_ -binding domain (IP_3_ BD; amino acids 224–604) of mouse IP_3_ receptor type I^17^ was amplified from mouse brain total RNA using RT-PCR. The IP_3_ BD was engineered by inserting XhoI and MluI restriction sites between residues Val435 and Ser436, then cloned into pRSET-A vector (Thermo Fisher Scientific, Waltham, MA, USA) at the BamHI/HindIII site. A DNA fragment encoding circularly permuted green fluorescent protein (cpGFP), derived from the prototype of the BGP Flashbody^18^, was digested with XhoI and MluI and inserted into the corresponding sites within the IP_3_ BD. To enhance cellular expression and solubility, the hyper-acidic region (amino acids 190–286) of the mouse amyloid precursor protein^19^ (NM_001198823.1) was fused to the N-terminus of IP_3_ BD. For mammalian expression, the final biosensors were subcloned into the pCAGGS vector^20^. Pink Flamindo^21^ plasmid was previously developed and R-GECO1^22^ plasmid was previously generated^23^ by DNA synthesis.To generate a plasmid for overexpression of the H1 receptor in HEK293T cells, a DNA fragment encoding *Homo sapiens* histamine receptor H1 was amplified using PCR from pH1R-P2A-mCherry-N1 (Addgene #84330), and cloned into the XhoI/NotI sites of CSII-EF-MCS (RIKEN BioResource Center RDB04378).

### 2.2 Biosensor optimization

To obtain a biosensor with better fluorescence intensity (FI) change (ΔF/F_0_) for IP_3_, the linker sequences at the N- and C-termini of cpGFP, which connect it to the IP_3_ -binding domain, were systematically varied by inserting or deleting residues to adjust the linker length over a range from - 6 to + 6 amino acids. All variants were expressed in *Escherichia coli* JM109(DE3) cells (Promega, Madison, WI, USA), and the ΔF/F_0_in crude lysates was analyzed for 10 μM IP_3_ . The variant showing the largest ΔF/F_0_was selected as the template for further mutagenesis.

To further improve the biosensor, site-directed saturated mutagenesis was performed on the linkers and adjacent amino acids using PCR with degenerate primers containing a mixture of NNK and MNN. Crude lysates prepared from approximately 50 colonies of transformed JM109(DE3) were analyzed in each round of molecular evolution. The variant exhibiting the largest ΔF/F_0_ was used as the template for subsequent iterative rounds of saturated mutagenesis. This molecular evolution process was continued until a variant with a response, which is F/F_0_ exceeding 3-fold was obtained, and this variant was named Green iPenguin 4.7. Subsequently, to generate biosensors with different affinities for IP_3_, additional saturated mutagenesis were performed to modulate the half-maximal effective concentration (EC_50_). This yielded three variants with distinct EC_50_ values, including Green iPenguin 0.67 and a negative control biosensor, Green iPenguin Nega.

### 2.4 Protein production and purification

Plasmids encoding Green iPenguins in the pREST-A vector were transformed into JM109(DE3). Transformed cells were cultured in 400 mL of LB medium supplemented with ampicillin at 20 °C for 4 days and collected by centrifugation. The pellets were resuspended in phosphate-buffered saline at pH 7.4 containing 0.5% (v/v) Triton X-100 (PBS-Triton), and the cells were disrupted using a high-pressure Cell Disruptor (Constant Systems Ltd., Daventry, UK). The lysate was centrifuged, and the supernatant was subjected to purification using TALON Metal Affinity Resin (TaKaRa Bio, Shiga, Japan). After washing the resin with PBS-Triton containing 10 mM imidazole, bound proteins were eluted with PBS-Triton containing 300 mM imidazole. To remove imidazole, the eluate was subjected to buffer exchange using a PD MidiTrap G-25 column (GE Healthcare, Chicago, IL, USA) pre-equilibrated with storage buffer containing 150 mM KCl, 50 mM HEPES-KOH, pH 7.4, 0.5% Triton X-100. The purified proteins were aliquoted and stored at -80 °C. Protein concentration was determined using the Bradford assay. A standard curve was generated using bovine serum albumin.

### 2.5 In vitro spectroscopy

Excitation and emission spectra of purified Green iPenguins were measured with a spectrophotometer F-2700 (Hitachi, Tokyo, Japan). Relative FI was calculated by dividing the FI in the presence of IP_3_ by that of the excitation and emission peaks around 490 and 510 nm in the absence of IP_3_, respectively. For dose-response curve,100 nM purified protein was incubated with 0.001–30 μM IP_3_ . EC_50_values were determined by four-parameter logistic curve fitting (eq1) using QtiPlot (IONDEV SRL, Bucharest, Romania).

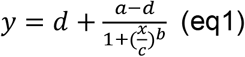

Specificity was evaluated by adding IP_3_ analogues at concentrations 10-fold higher than their EC_50_ values.

The absorption spectra of 20 μM Green iPenguin proteins were measured by a spectrophotometer V-730 BIO (JASCO Corporation, Tokyo, Japan) in the presence or absence of 30 μM IP_3_.

### 2.6 Cell culture and transfection

HeLa, HEK293T, and MCF7 cells were cultured in Dulbecco’s modified Eagle’s medium (high glucose) with L-glutamine, sodium pyruvate, penicillin, and streptomycin (FUJIFILM Wako Pure Chemical Corporation, Osaka, Japan), and 10% (v/v) heat-inactivated fetal bovine serum (GE Healthcare, Chicago, IL, USA). These cells were seeded in 35 mm glass-bottomed dishes at 37 °C with 5% CO_2_ for live-cell imaging. Plasmids of Green iPenguins were transfected into these cells using Lipofectamine 3000 Transfection Reagent (Thermo Fisher Scientific, Massachusetts, MA, USA) following the manufacturer’s instructions. After cells were cultured at 37 °C for 4 to 6 hours, the medium was replaced. For chromophore maturation, cells were cultured at 30 °C for at least 48 hours.

### 2.7 Live-cell imaging

Live-cell imaging was performed using an inverted microscope IX70 (Olympus, Tokyo, Japan) equipped with an oil-immersion ×40 objective lens, UApo/N 340, NA = 1.35, a cooled CCD camera, CoolSNAP HQ2 (Photometrics, Tucson, AZ, USA), and a mercury lamp. For GFP imaging, U-MGFPHQ, an excitation filter of 460–480 nm, an emission filter of 495–540 nm, and a dichroic mirror of 485 nm (Olympus, Tokyo, Japan) were used. For dual-color imaging, the DA/FI/TR-3x3M-C Sedat filter set (OPTO-LINE, Inc., Saitama, Japan), the green channel with an excitation filter of 480/17 nm and an emission filter of 520/28 nm, the red channel with an excitation filter of 556/20 nm and an emission filter of 617/73 nm, and the triple-band dichroic mirror (403/497/574 nm) was used. The filter exchange was performed using an HF110 high-speed filter wheel (Prior Scientific, Cambridge, UK). Fluorescence images were acquired every 5 seconds using Metafluor software (Molecular Devices, San Jose, CA, USA). For IP_3_ -Ca^2+^ hierarchy analysis, time-lapse imaging was performed at the maximum acquisition rate, approximately 1 second per frame, with slight variations in frame intervals due to channel switching and acquisition delays. The FI was measured by manually surrounding each cell with a region of interest (ROI) using ImageJ software (NIH, Bethesda, MD, USA). Background FI was subtracted using a cell-free area. Relative FI was calculated by normalizing to the average FI during a 30 or 60 second baseline period prior to stimulus addition.

## 3 Results

### 3.1 Generation of Green iPenguins

We generated a genetically encoded IP_3_ biosensor by inserting the circularly permuted green fluorescence protein (cpGFP) into IP_3_ -binding domain (IP_3_ BD), which is amino acids 224–604 of mouse IP_3_ receptor type 1. To identify the best insertion position for generating the biosensor, thirteen variants with different insertion positions within the IP_3_ BD were constructed (Fig. S1A). To examine the response against IP_3_, these variants were expressed in *E. coli*, and the fluorescence intensity (FI) of the cell lysates was measured with or without 10 μM IP_3_ . The variant with cpGFP inserted between V435 and S436, which is position 10 displayed the largest response, approximately 1.04-fold. To further enhance the response, we adjusted the linker lengths between cpGFP and IP_3_ BD by deletion or addition of amino acids. The linker was based on the leucine zipper, with the expectation that its strong α-helical structure would improve the response similarly to previous studies. The variant with N ± 0 aa and C + 1 aa (Fig. S1B), showing the largest response of approximately 1.1-fold, was further improved by saturated mutagenesis using overlap PCR, targeting amino acids within and around the linkers. After 12 cycles of saturated mutagenesis, we obtained the biosensor with more than 3-fold response in the presence of 10 μM IP_3_, which was named Green iPenguin 4.7 (Fig. S2). The biosensor showed an EC_50_ slightly outside the physiological IP_3_ concentration range, from 40 nM to 1.8 μM^24^. Therefore, we performed additional mutagenesis using a physiological IP_3_ concentration of 1 μM for screening, and obtained Green iPenguin 0.67, which responds within the physiological range. Two further rounds of mutagenesis were performed to generate the negative-control biosensor, Green iPenguin Nega. As described above, we successfully generated three green IP_3_ biosensors (Fig. 1A). The numerical part of each name reflects the EC_50_ value, which is determined in the following section (Fig. 1B).

**Figure 1.**
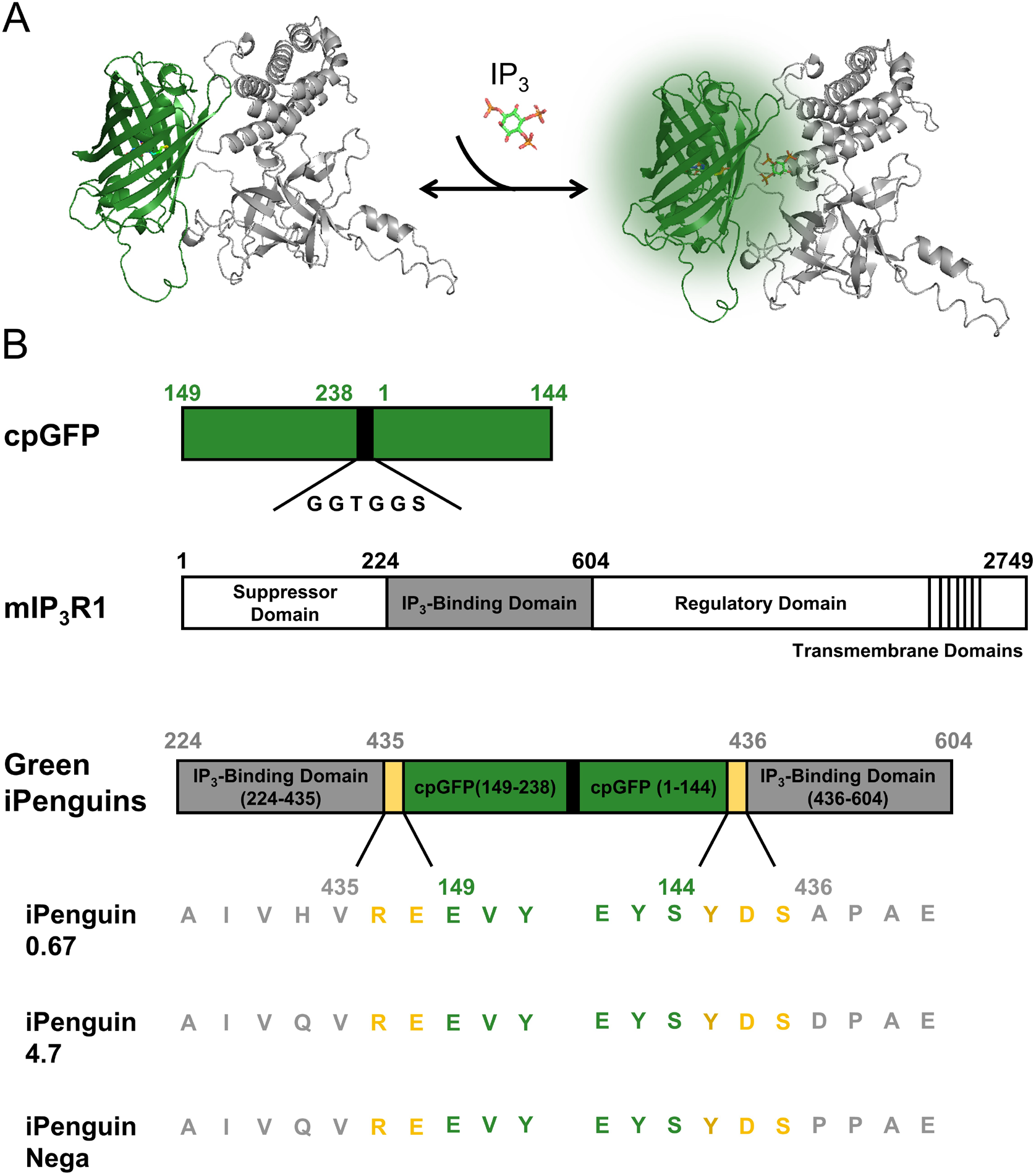
Schematic drawing of Green iPenguins. (A) Schematic 3D structures of Green iPenguins unbound (left) and bound (right) to IP_3_ . The structures were predicted by AlphaFold2. (B) Diagrams for cpGFP and mouse IP_3_ receptor type1, and amino acid sequence near linker regions showing differences among Green iPenguin 0.67, 4.7, and Nega.

### 3.2 Characterization of Green iPenguins

To examine the spectral property and dose-response relationships, we purified the protein of Green iPenguins produced in *E. coli*. These Green iPenguins have excitation peaks at around 490 nm and emission peaks at around 510 nm, which are similar to GFP^25^. The response of Green iPenguin 0.67 and 4.7 was 4.5- and 5.4-fold, respectively, whereas that of Green iPenguin Nega was less than 1.1-fold. (Fig. 2A– C). The absorption spectra of Green iPenguin 0.67 and 4.7 showed a decrease in the absorption peak around 400 nm and an increase around 500 nm in the presence of IP_3_, respectively, whereas Green iPenguin Nega did not show a distinguishable change (Fig. S3A–C). These results suggest that the change in FI upon IP_3_ addition was caused by an alteration to the protonated/deprotonated state of the chromophore, which is a general mechanism^23^ underlying single fluorescent protein (FP)-based biosensors. When we investigated the response for various concentration of IP_3_, FI was increased in a dose-dependent manner (Fig. 2D). The biosensors were named Green iPenguin 0.67 and 4.7 based on their EC_50_ values of 0.67 and 4.7 μM, respectively. Then, to evaluate the specificity of Green iPenguins for IP_3_, we added structurally related inositol phosphates (1,3,4-IP_3_, 1,4-IP_2_, 1,3,4,5-IP_4_) as well as Gq– phospholipase C (PLC) signaling molecules (PIP_2_ and DAG) at 10 times the concentration of EC_50_ against IP_3_ for each biosensor, which is 6.7 and 47 μM. For Green iPenguin Nega, IP_3_ was added at 47 μM, the highest test concentration, because its EC_50_ value was not able to be determined. The Green iPenguin 0.67 and 4.7 displayed high specificity for IP_3_, with only minor responses to 1,3,4,5-IP_4_ and PIP_2_, and Green iPenguin Nega didn’t show distinguishable response (Fig. 2E). Taken together, these results suggest that Green iPenguins respond specifically to IP_3_ and are potentially suitable for visualizing IP_3_ dynamics in living cells.

**Figure 2.**
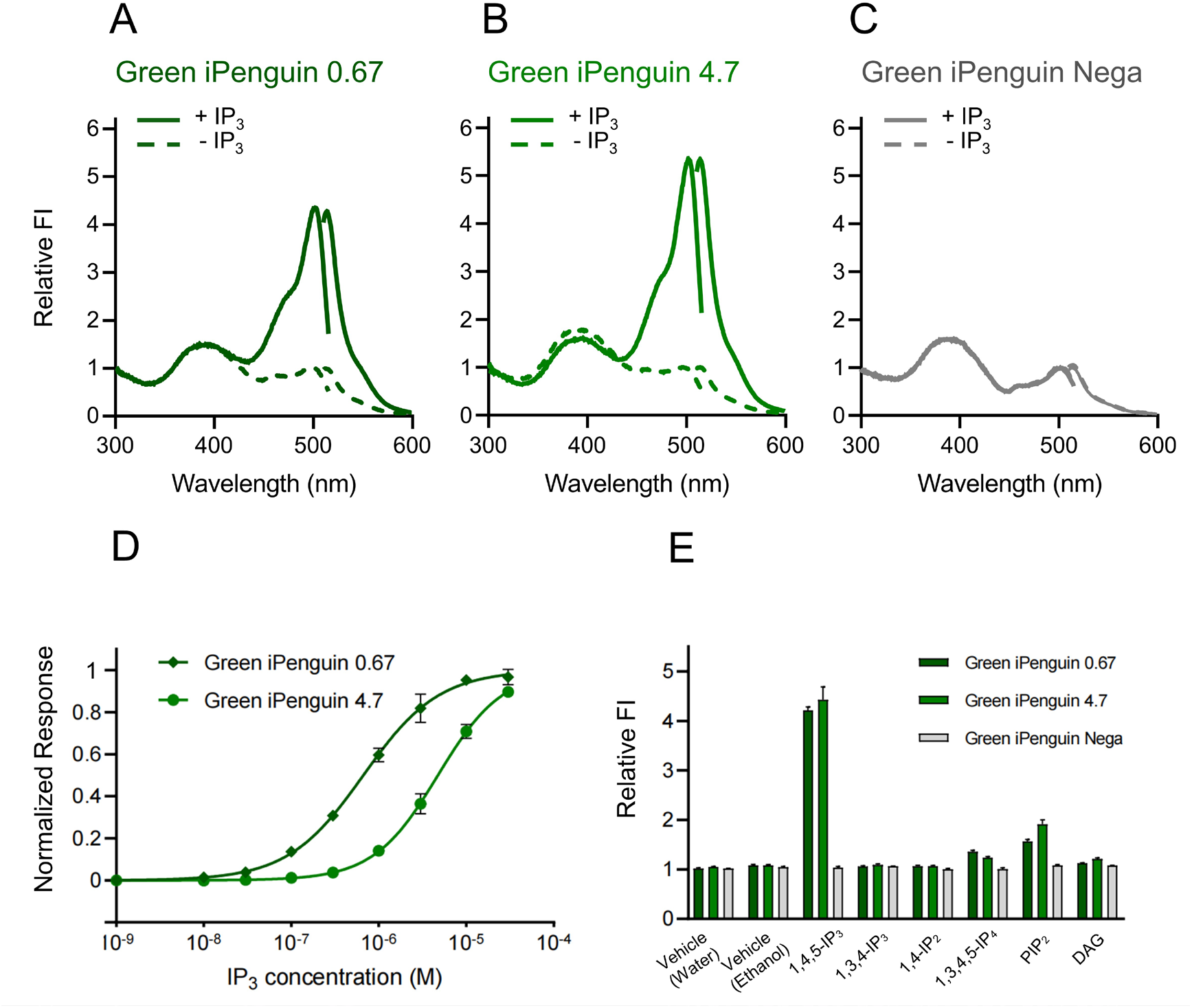
Spectral properties of Green iPenguins. (A–C) Excitation and emission spectra of Green iPenguin 0.67, Green iPenguin 4.7, and Green iPenguin Nega in the presence (solid line) and absence (dashed line) of 30 μM IP_3_ . (D) Dose-response curve of Green iPenguins against IP_3_ . (E) Specificity of purified Green iPenguins for IP_3_ related molecules. The data represent the means ± standard deviation (SD) of three independent measurements performed using the same batch of samples.

### 3.3 Green iPenguin visualizes IP_3_ dynamics in living cells

To investigate the applicability of Green iPenguins for live-cell imaging, we expressed Green iPenguin 0.67 and Nega in HEK293T cells and monitored FI after application of various ligands. Green iPenguin 0.67 was employed as its EC_50_ for is in the physiological range of IP_3_ . Upon application of 100 μM carbachol^26^ (Cch) for as an agonist of the muscarinic acetylcholine receptor M3, or 10 μM ATP^27^ as an agonist of the P2Y_2_ purinergic receptor, the FI of Green iPenguin 0.67 increased, while Green iPenguin Nega showed only minor changes (Fig. 3A–C). Having validated Green iPenguins for monitoring endogenous receptor-mediated signaling, we next examined their performance with exogenously introduced receptors to demonstrate the potential applicability of the biosensors to future functional analyses of diverse GPCRs, including orphan GPCRs, and their mediated signaling pathways. In HEK293T cells stably expressing human H1 receptor (H1R) by lentiviral transduction, application of histamine induced around a 6-fold increase in FI of Green iPenguin 0.67, but not in Green iPenguin Nega, and response was not induced in HEK293T cells without exogenous H1R expression (Fig. 3D). Treatment with 10 μM U73122^28^, a PLC inhibitor, abolished the agonist-induced FI increases, confirming that all observed responses were PLC-dependent (Fig. 3E–G). Taken together, these results suggest that the Green iPenguin is applicable to monitor IP_3_ dynamics across a variety of GPCR-mediated signaling pathways, even when the receptors are introduced exogenously.

**Figure 3.**
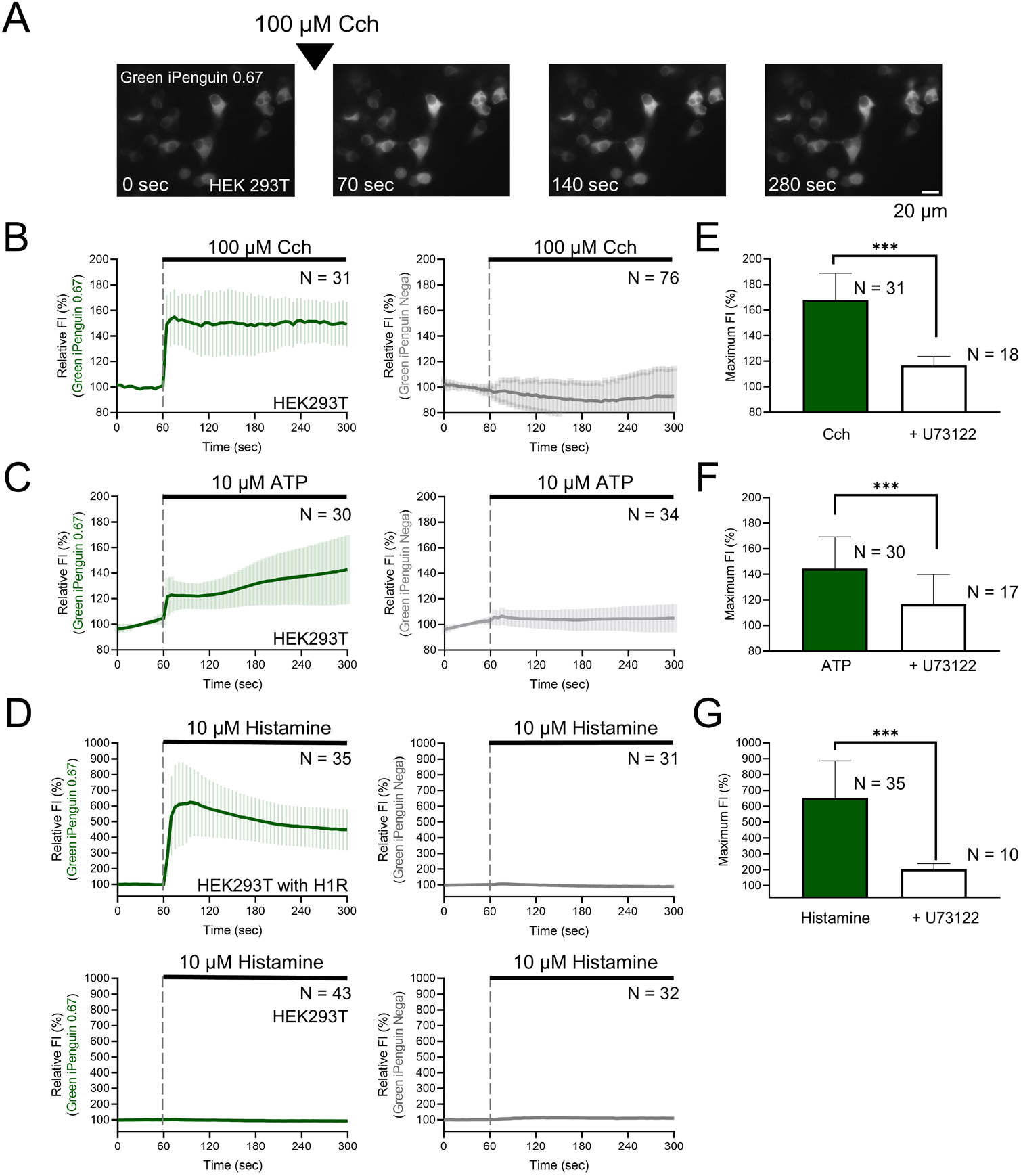
Live-cell imaging using Green iPenguins. (A) Representative sequential images of Green iPenguin 0.67 expressing HEK293T cells during the application of 100 μM Cch. The triangle (▼) indicates the time point of agonist application. (B, C) Time courses of FI in Green iPenguin 0.67/Nega expressing HEK293T cells during the application of 100 μM Cch or 10 μM ATP. (D) Time courses of FI in Green iPenguin 0.67/Nega expressing in HEK293T cells with exogenously introduced H1R or in wild-type HEK293T cells during the application of 10 μM histamine. (E–G) The maximum FI during the application of ATP, Cch, or histamine with 10 μM U73122 treatment. The numbers in bar graphs represent the number of cells analyzed from more than three independent experiments. The data represent the means ± SD. ***p < 0.001 by Student’s t-test.

### 3.4 Dual-color imaging for visualizing cooperative and hierarchical dynamics

The second messengers act cooperatively and hierarchically to regulate diverse cellular processes, and elucidating their interplay requires simultaneous monitoring within the same cell. We co-expressed Green iPenguin 0.67 and Pink Flamindo^21^, a red cAMP biosensor, in MCF7 cells, enabling simultaneous monitoring of cAMP and IP_3_, which has been difficult to achieve with previous biosensors. When 100 μM adrenaline^29^ for an agonist of both α1- and β-adrenergic receptors was applied, the FI of both biosensors increased (Fig. 4A–C). Next, we co-expressed Green iPenguin 0.67 and the red fluorescent Ca^2+^ biosensor R-GECO1^22^ in HeLa cells, and monitored IP_3_ and Ca^2+^ dynamics, which had rarely been performed because it required a complex microscope setup for combination of a genetically encoded FRET-based IP_3_ biosensor and a chemical Ca^2+^ dye. After application of 10 μM histamine, the FI of both biosensors increased (Fig. 4D–F). These results suggest that Green iPenguin 0.67 is well suited for dual-color imaging and can be used in combination with biosensors of different colors.

**Figure 4.**
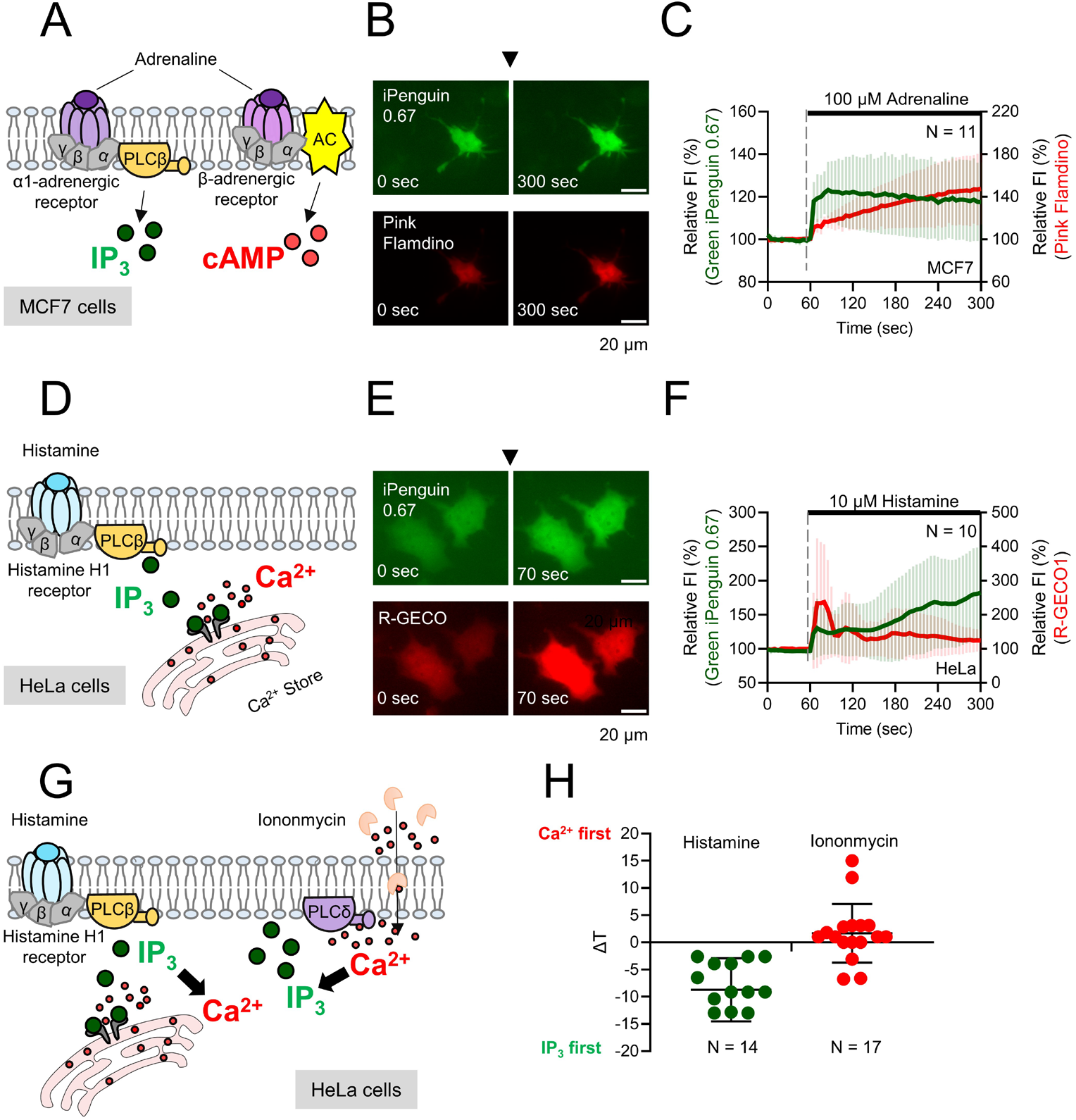
Dual-color imaging for visualizing the IP_3_ with cAMP or Ca^2+^. (A–C) Sequential images and time course of FI of Green iPenguin 0.67 and red genetically encoded cAMP biosensor, Pink Flamindo co-expressing MCF7 cells during the application of 100 μM adrenaline. (D–F) Sequential images and time course of FI of Green iPenguin 0.67 and red genetically encoded Ca^2+^ biosensor, R-GECO1 co-expressing HeLa cells during the application of 10 μM histamine. (G, H) Comparison of the onset time difference between IP_3_ and Ca^2+^ signals during the application of histamine and ionomycin. The triangle (▼) indicates the time point of agonist application. The numbers in graphs represent the number of cells analyzed from more than three independent experiments. Data are presented as mean ± SD.

However, this dual-color imaging does not resolve the temporal order between the signals. Applying the same approach with higher temporal resolution enables a more detailed analysis of the hierarchical dynamics of IP_3_ and Ca^2+^ and allows direct visualization of IP_3_ preceding Ca^2+^ mobilization in the Gq–PLC signaling pathway. To examine the hierarchical relationship between IP_3_ and Ca^2+^ dynamics, we performed dual-color imaging with high temporal resolution with histamine or ionomycin. Histamine activates Gq-coupled receptors, triggering PLCβ activation, IP_3_ production, and subsequent Ca^2+^ mobilization from intracellular stores. On the other hand, ionomycin acts as a Ca^2+^ ionophore that directly elevates cytosolic Ca^2+^ levels, thereby inducing Ca^2+^-dependent IP_3_ production through PLCδ. We defined the onset as the first frame in which the signal exceeded the baseline (mean + 3 SD) and continued to rise for at least five consecutive frames. Positive and negative values reflect earlier Ca^2+^ and IP_3_ elevations, respectively. After histamine application, IP_3_ elevation consistently preceded Ca^2+^ increases. In contrast, following ionomycin application, Ca^2+^ increases preceded IP_3_ elevation in the majority of cells (Fig. 4G, H). Taken together, these results suggest that Green iPenguin 0.67 enables visualization of hierarchical relationships between IP_3_ and Ca^2+^ and can be extended to reveal hierarchical relationships between IP_3_ and other signaling molecules.

## 4. Discussion

Green iPenguins, single FP-based IP_3_ biosensors, were developed through iterative molecular design combined with directed evolution. Based on the success of FRET-based biosensors, we employed the IP_3_ BD derived from mouse IP_3_ receptor type 1 and inserted cpFP into IP_3_ BD. This design strategy enables ligand-induced conformational changes in the binding domain to be transmitted to the chromophore environment of cpFP, thereby modulating fluorescence intensity^30^.This strategy was chosen because our previous attempts to insert IP_3_ BD into GFP, as used in our earlier studies^31^, were unsuccessful, likely because of the insertion of relatively large size of IP_3_ BD. However, a known limitation of biosensors inserting cpFP is that the engineering the binding domain, such as the insertions or amino acid substitutions, may alter binding affinity and specificity. Consistent with our expectation, the EC_50_ value of Green iPenguin 0.67 was 0.67 μM, which is higher than the reported *K*_D_ value around 2 nM of the isolated IP_3_ BD, reflecting a reduced binding affinity. Nevertheless, this EC_50_ value remains within the physiological concentration range such as 40 nM to 1.8 μM^24^, supporting the applicability of Green iPenguin 0.67 for monitoring intracellular IP_3_ dynamics. On the other hand, specificity analyses revealed that Green iPenguins responded to 1,4,5-IP_3_ strongly, with no significant responses to 1,3,4-IP_3_, 1,4-IP_2_, or DAG, and only weak responses to 1,3,4,5-IP_4_ and PIP_2_. A weak response to 1,3,4,5-IP_4_ has also been reported for previously developed FRET-based IP_3_ biosensors. Importantly, the response to 1,3,4,5-IP_4_ was minimal, and PIP_2_ is spatially restricted to the plasma membrane with limited diffusional mobility. Therefore, their contributions to cytosolic signals are expected to be negligible under physiological conditions.

To further examine the utility of Green iPenguin in live-cell imaging, we expressed Green iPenguin in HEK293T, and monitored the IP_3_ dynamics across different GPCR-mediated signaling pathways. We found that application of either ATP or Cch induced responses of less than 2-fold. This was markedly lower than the approximately 6-fold response observed upon histamine stimulation in cells in which the histamine H1 receptor was exogenously introduced, which was slightly higher than that of the purified protein. These differences are thought to arise from distinct causes. One is that exogenously introduced receptors exhibit higher expression levels than under endogenous conditions, leading to increased IP_3_ production compared with the physiologically small changes in IP_3_ . Another is that intracellular expression of the biosensor avoids the physical damage associated with protein purification. Importantly, despite these differences, these results indicate that Green iPenguin can monitor IP_3_ dynamics both through endogenously expressed and exogenously introduced receptors, rendering it suitable for applications such as investigating GPCR crosstalk and screening for orphan GPCR ligands.

In the hierarchy analysis of IP_3_ and Ca^2+^ using dual-color imaging, we observed that the time interval between IP_3_ and Ca^2+^ signals upon histamine application was longer than that observed between Ca^2+^ and IP_3_ signals upon ionomycin application (Fig. 4H). This difference likely arises because histamine induces IP_3_ production at the plasma membrane, requiring diffusion of IP_3_ to the endoplasmic reticulum before Ca^2+^ mobilization occurs, whereas ionomycin directly increases cytosolic Ca^2+^, leading to rapid activation of Ca^2+^-dependent PLC and IP_3_ production. Thus, the relative timing of IP_3_ and Ca^2+^ signals reflects differences in the temporal order of IP_3_ and Ca^2+^ signaling, including the location where the signaling molecules originate. This approach will allow the precise analysis of both the site of signaling molecule production and their diffusion dynamics within the cell, offering powerful insights into intracellular signaling. Considering that GPCRs have been reported in multiple subcellular compartments^32^, such studies are expected to open exciting avenues for future research.

## ASSOCIATED CONTENT

## AUTHOR INFORMATION

### Author contributions

T.K. conceived the study. L.T. and T.K. designed the experiments and wrote the manuscript. L.T., K.Y. and F.Z. performed the experiments and analyzed the data. B.Z.,T.Y. and T.T. designed the experiments and edited the manuscript. All authors approved the final version of the manuscript.

## DATA AVAILABILITY

Raw data are available from the corresponding author upon reasonable request.

