## Supporting Information for "Green genetically encoded IP_3_ biosensor for hierarchical analysis of its signaling network"

**Table of Contents**

**Figure S1.** Schematic drawing for the screening processes of Green iPenguins.
**Figure S2.** Diagram for the site-directed saturated mutagenesis.
**Figure S3.** Absorption spectra of Green iPenguins.

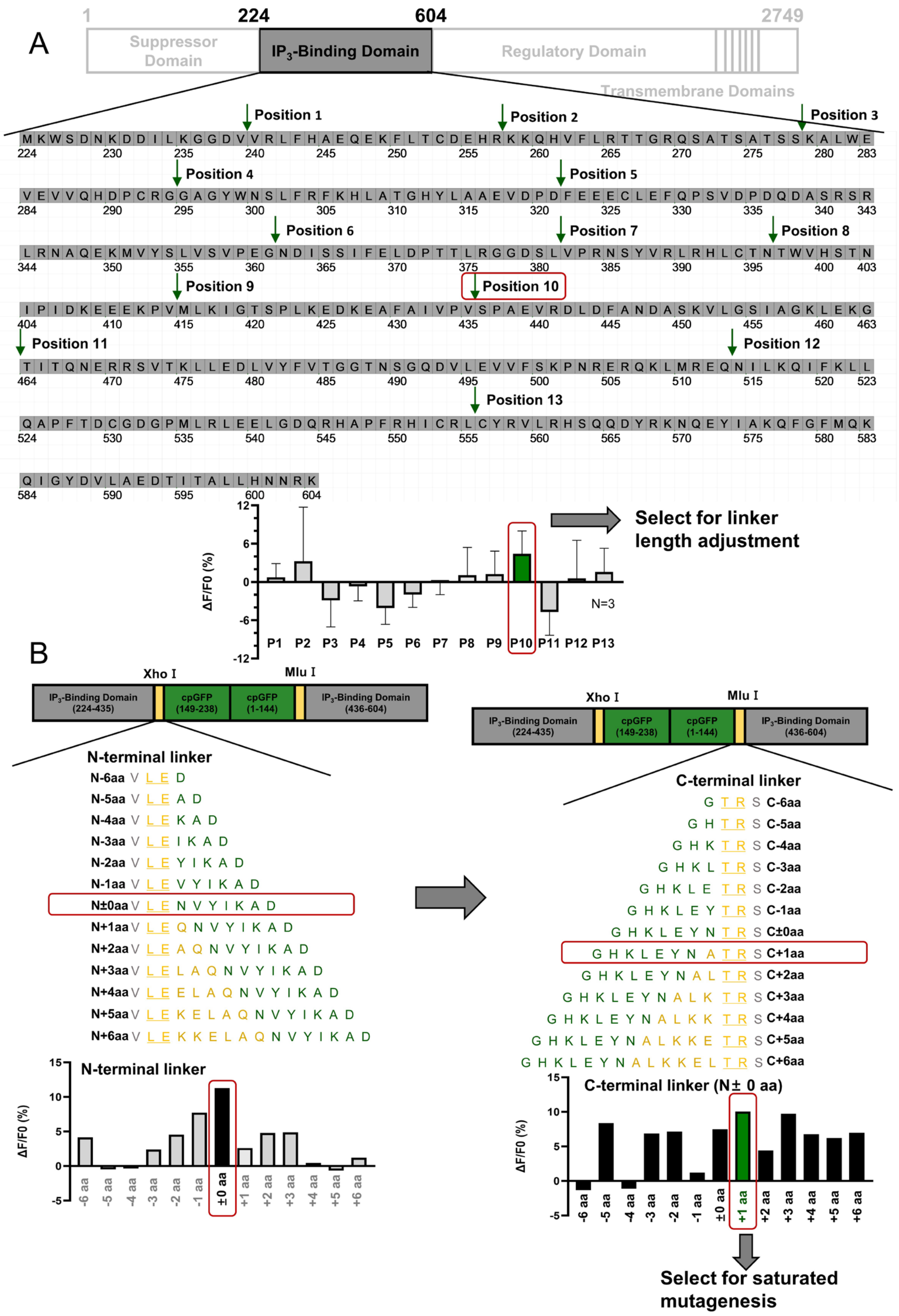


**Figure S1.** Schematic drawing for the screening processes of Green iPenguins. (A) Amino acid sequence of the IP$_3$ binding domain with inserted position of cpGFP, and ΔF/F$_0$ of the variants with different insertion positions upon 10 μM IP$_3$. (B) Diagrams of the Green iPenguins with different linker lengths at the N- and C-termini of cpGFP, and ΔF/F$_0$ of the variants with different linker lengths upon 10 μM IP$_3$.

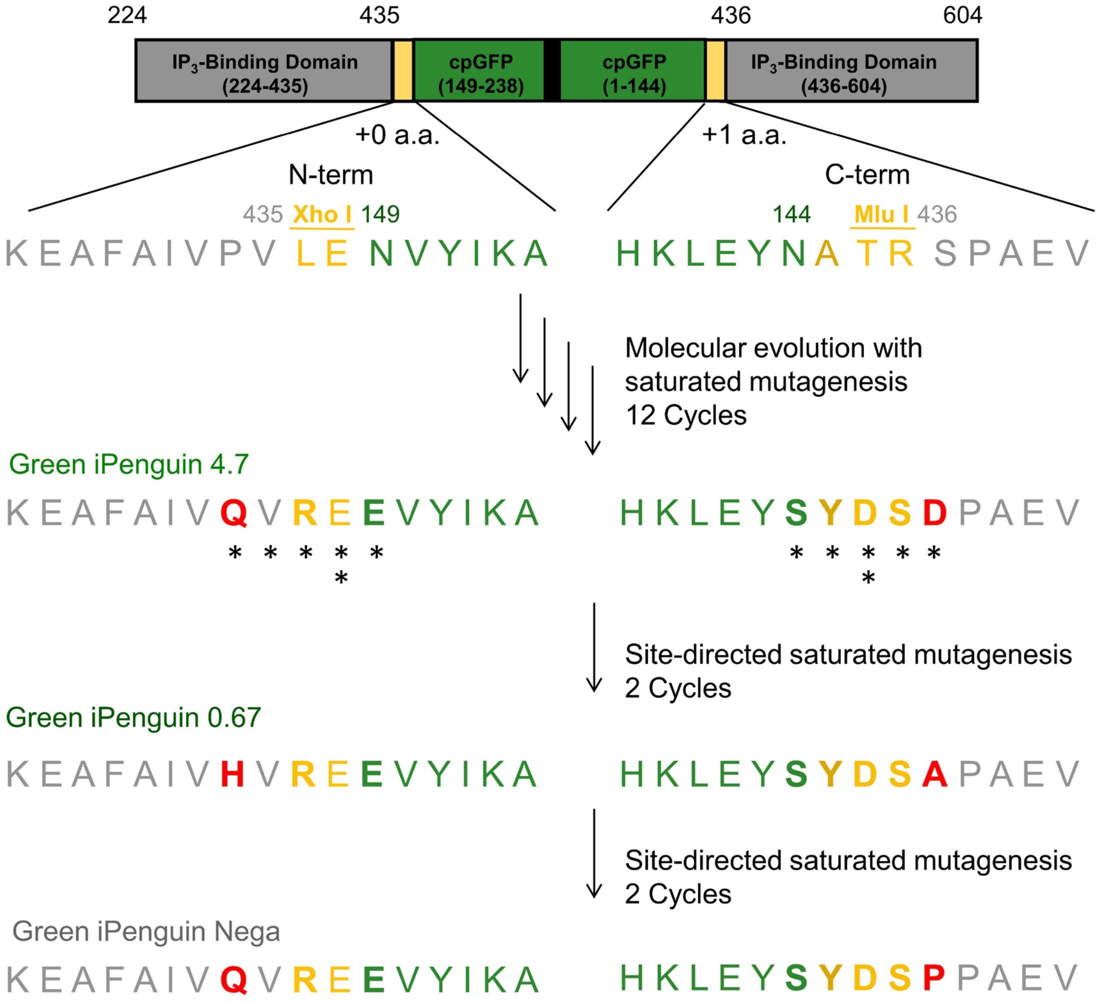


**Figure S2.** Diagram for the site-directed saturated mutagenesis. The saturated mutagenesis was carried out on the linker-optimized variant (N ± 0 aa, C + 1 aa). Asterisks indicate the residues targeted for mutagenesis, and the number of asterisks represents the number of mutagenesis attempts. Substituted amino acids are shown in bold. The amino acids differences among the three biosensors with distinct $EC_{50}$ are indicated in red.

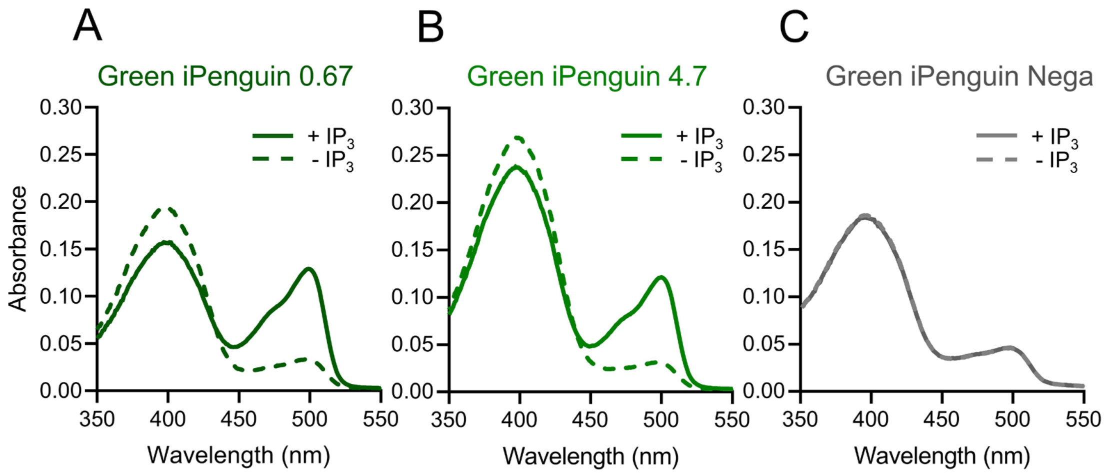


**Figure S3.** Absorption spectra of Green iPenguins. (A–C) Absorption spectra of Green iPenguin 0.67, Green iPenguin 4.7, and Green iPenguin Nega in the presence (solid line) and absence (dashed line) of 30 µM $IP_3$.
